# Human Erbb2-induced Erk Activity Robustly Stimulates Cycling and Functional Remodeling of Rat and Human Cardiomyocytes

**DOI:** 10.1101/2020.12.15.422603

**Authors:** Nicholas Strash, Sophia DeLuca, Geovanni L. Janer Carattini, Soon Chul Heo, Ryne Gorsuch, Nenad Bursac

**Affiliations:** Department of Cell Biology, Duke University, Durham NC; Department of Biomedical Engineering, Duke University, Durham NC

## Abstract

Multiple mitogenic pathways capable of promoting mammalian cardiomyocyte (CM) proliferation have been identified as potential candidates for functional heart repair following myocardial infarction. However, it is unclear whether the effects of these mitogens are species-specific and how they directly compare in the same cardiac setting. Here, we examined how CM-specific lentiviral expression of various candidate mitogens affects human induced pluripotent stem cell-derived cardiomyocytes (hiPSC-CMs) and neonatal rat ventricular myocytes (NRVMs) *in vitro*. In 2D-cultured CMs from both species, and in highly mature 3D-engineered cardiac tissues generated from NRVMs, a constitutively-active mutant form of the human gene Erbb2 (cahErbb2) was the most potent tested mitogen. Persistent expression of cahErbb2 induced CM proliferation, sarcomere loss, and remodeling of tissue structure and function, which were attenuated by small molecule inhibitors of Erk signaling. These results suggest transient activation of Erbb2/Erk axis in cardiomyocytes as a potential strategy for regenerative heart repair.

## Introduction

The adult mammalian heart is composed primarily of post-mitotic CMs (Bergmann et al., 2009; Mollova et al., 2013). Due to an apparent lack of a resident stem cell population (van Berlo et al., 2014), the heart is unable to replace lost CMs following myocardial infarction (MI), and instead undergoes fibroblast-mediated scarring, resulting in a decline in cardiac function. One potential approach to regenerate heart tissue following MI is to stimulate the proliferation of survived CMs by a gene therapy targeting cell cycle regulation (Leach & Martin, 2018; Lin et al., 2014; Mohamed et al., 2018; Tzahor & Poss, 2017). While inducing adult CM proliferation is only the first step towards achieving efficient cardiac repair post-MI, it is arguably the most challenging and important one. Which of the proposed candidate genes and pathways would optimally induce human CM proliferation to eventually restore cardiac tissue function remains to be determined.

Several approaches involving increased expression of cell cycle-related proteins and modulation of the Hippo, neuregulin (Nrg1), or Wnt signaling pathways have been shown to generate robust proliferative responses in rodent and zebrafish CMs. Overexpression of cyclin D2 alone or a combination of cyclin-dependent kinase 1 (CDK1), CDK4, cyclin B1, and cyclin D1 promoted cell cycle activation in post-mitotic mouse, rat, and human CMs *in vitro* and in mouse hearts *in vivo* (C. Fan et al., 2019; Mohamed et al., 2018; Zhu, Zhao, Mattapally, Chen, & Zhang, 2017). Hippo pathway modulation via overexpression of constitutively active (ca) Yap variants (incapable of being phosphorylated and degraded) led to robust proliferative responses *in vitro* and *in vivo* through chromatin alterations surrounding proliferation-inducing genes (Byun et al., 2019; Monroe et al., 2019). The mitogen Nrg1 induced CM proliferation in zebrafish through activation of its co-receptor, Erbb2 (Bersell, Arab, Haring, & Kuhn, 2009; Gemberling, Karra, Dickson, & Poss, 2015), and overexpression of caErbb2 promoted CM proliferation in adult mice by inducing their de-differentiation (D’Uva et al., 2015), possibly via Yap activation (Aharonov et al., 2020). Wnt stimulation by small molecule inhibition of GSK3β (Buikema et al., 2020; Y. Fan et al., 2018; Richard J. Mills et al., 2019) or induction of β-catenin release from the cell membrane (Y. Fan et al., 2018) were shown to induce proliferation in human CMs *in vitro*. Yet, there have been no studies that directly compared mitogenic capacity of these different pathways in the same CM preparation.

In this report, we sought to directly compare manipulations of different signaling pathways for their ability to induce cell cycle activation and proliferation in CMs from various species and maturation levels, including 2D monolayer cultures of human induced pluripotent stem cell-derived CMs (hiPSC-CMs (I. Y. Shadrin, Khodabukus, A. & Bursac, N, 2016; Zhang et al., 2013)) and neonatal rat ventricular myocytes (NRVMs), as well as functional 3D engineered cardiac tissues (“cardiobundles”) made from NRVMs (Helfer & Bursac, 2020; C. P. Jackman, Carlson, & Bursac, 2016). We then probed the mechanisms underlying the observed proliferative responses. We confirmed that Erk signaling is a major contributor to Erbb2-induced cell cycle activation (D’Uva et al., 2015), and found that the human caErbb2 ortholog (cahErbb2) induces more potent proliferative effects in hiPSC-CMs and NRVMs than rat caErbb2 (carErbb2) (Aharonov et al., 2020; D’Uva et al., 2015), which was unable to activate Erk signaling in human CMs.

## Results

### Lentiviral Expression of Mitogens Promotes hiPSC-CM Proliferation without Inducing Apoptosis

Lentiviral vectors (LVs) were designed using the MHCK7 promoter (Salva et al., 2007) to drive muscle-specific expression of mitogens together with an mCherry reporter and applied to transduce hiPSC-CMs and NRVMs (Figure 1 – figure supplement 1). We used a flow cytometry strategy (Figure 1 – figure supplement 2) validated using an established hiPSC-CM mitogen (GSK3 inhibitor, CHIR99021) (Buikema et al., 2020; Richard J. Mills et al., 2019) to determine whether a LV-induced mitogen expression resulted in cell cycle activation or apoptosis specifically in mCherry-labelled CMs. Compared to LV-driven expression of mCherry only, hiPSC-CMs transduced with LVs (Figure 1A) coding for caCtnnb1 (ca β-catenin), Ccnd2 (Cyclin D2), carErbb2, or cahErbb2, but not caYap8SA exhibited significantly higher incorporation of 5-ethynyl-2′-deoxyuridine (EdU), indicating greater DNA synthesis (Figure 1B). Increased numbers of CMs expressing phosphorylated histone H3 (H3P), an indicator of the mitotic (M) phase of the cell cycle, were also found with cahErbb2 and Ccnd2 transduction (Figure 1C). Furthermore, we observed comparable percentage of diploid and polyploid hiPSC-CMs between control and LV-treated total or EdU^+^ CMs (Figure 1D,E). Collectively, these results showed that hiPSC-CMs with LV-mediated expression of cahErbb2 and Ccnd2 were induced to enter the DNA synthesis and mitotic phases of the cell cycle at a higher rate than control hiPSC-CMs, and continued to undergo succesful cytokinesis. Additionally, from cleaved caspase-3 (Cc3) analysis, transduced mitogens did not increase CM apoptosis, with caYap8SA having an anti-apoptotic effect (Figure 1F). In contrast, both increased DNA synthesis and mitosis as well as pro-apoptotic effects were found with the application of CHIR99021 (Figure 1 – figure supplement 1). Together, increased EdU incorporation and no change in CM polyploidy or apoptosis upon transduction with caCtnnb1, Ccnd2, carErbb2, or cahErbb2 LVs indicated induced CM proliferation.

**Figure 1.**
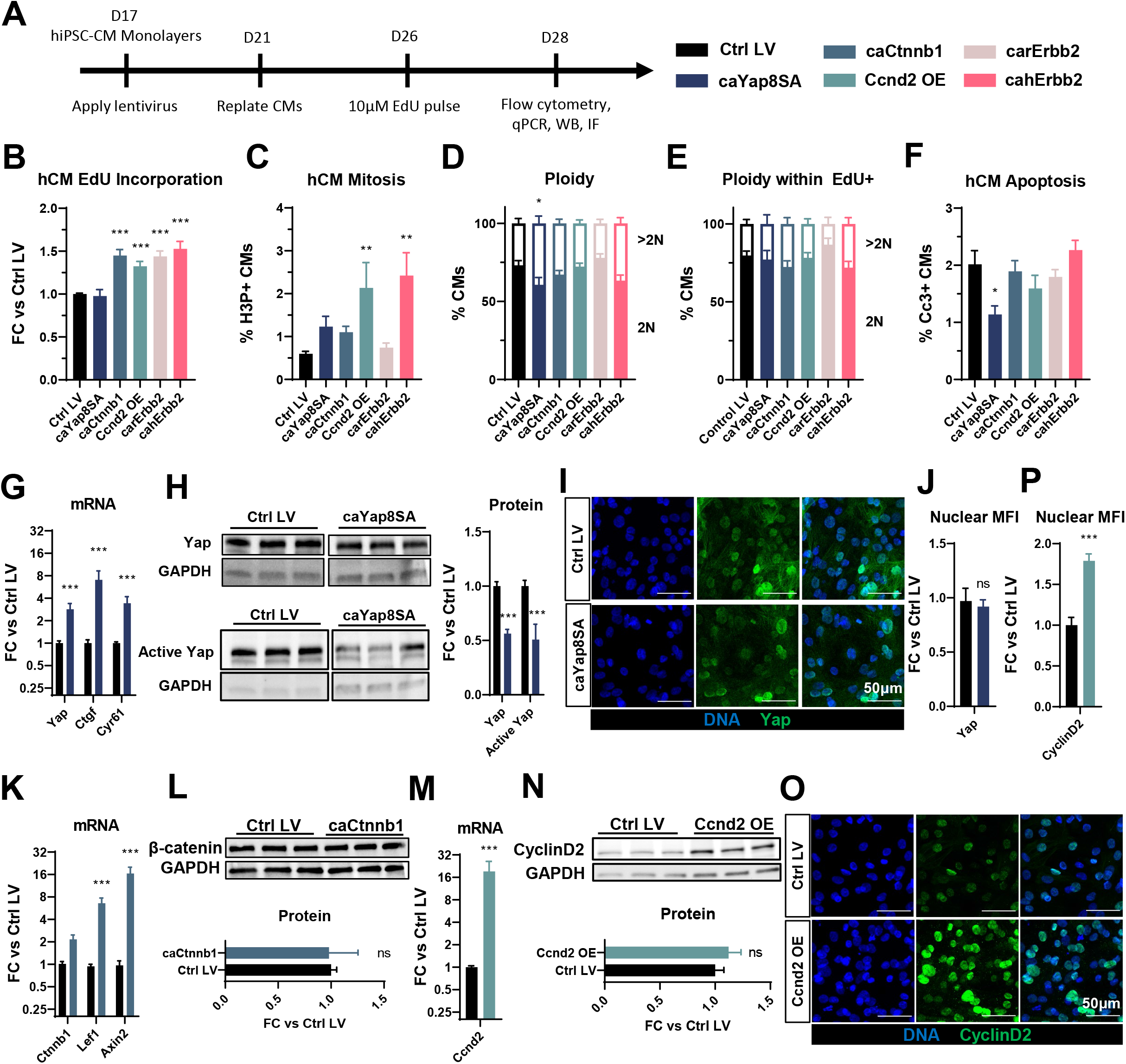
LV-delivered Mitogens Drive hiPSC-CM Proliferation in Monolayers. (A) Schematic of experimental design in hiPSC-CM monolayers. (B-F) Flow cytometry analysis of mCherry^+^ hiPSC-CMs showing (B) fold-change (FC) in EdU incorporation relative to control LV-transduced hiPSC-CMs, (C) percentage of H3P^+^ Ms, (D,E) percentage of 2N and >2N cells in all CMs (D) and EdU^+^ CMs (E), and (F) percentage of apoptotic Cc3^+^ CMs. (G,H) Analysis of relative (G) gene expression of *Yap* and its targets *Ctgf* and *Cyr61*, and (H) total and active Yap protein abundance in caYap8SA-transduced vs. control hiPSC-CMs. (I,J) Representative imunnostaining images (I) and quantified nuclear Mean Fluorescence Intensity (MFI, J) of YAP in caYap8SA-transduced vs. control hiPSC-CMs. (K-L) Analysis of relative (K) expression of *Ctnnb1* and Wnt-signaling genes *Lef1* and *Axin2* and (L) Ctnnb1 protein abundance in caCtnnb1-transduced vs. control hiPSC-CMs. (M,N) Analysis of relative Ccnd2 (M) gene and (N) protein expression and (O,P) Representative immunostaining images (O) and quantified nuclear MFI (P) of Ccnd2 in Ccnd2-transduced vs. control hiPSC-CMs. Data: mean ± SEM (*p < 0.05, **p < 0.01, ***p < 0.001 vs. Ctrl LV). See Table 1 for sample numbers and complete statistical information for all Figures.

### Lentiviral Expression of Mitogens in hiPSC-CMs Activates Negative Feedback Loops

We then assessed molecular effects of LV transduction and found that caYap8SA-transduced hiPSC-CMs exhibited significantly increased gene expression of *Yap* and its downstream targets *Ctgf* and *Cyr61* (Figure 1G). However, the expression of the total and active, non-phosphorylated Yap protein was significantly decreased (Figure 1H). Immunostainng of transduced hiPSC-CMs further revealed that the nuclear abundance of active Yap was unchanged by caYap8SA expression (Figure 1I,J). Because caYap8SA cannot be phosphorylated and degraded, these studies imply the existence of a negative feedback loop that degraded endogenous Yap protein to compensate for caYap8SA overexpression. This compensatory mechanism was likely responsible for the lack of mitogenic effects in caYap8SA transduced hiPSC-CMs. We further assessed caCtnnb1-transduced hiPSC-CMs and found that the gene expression of *Ctnnb1* was unaltered, but noted an increased expression of *Lef1* and *Axin2* (Figure 1K). This suggested increased downstream Wnt pathway activity that may have resulted in a negative feedback loop to suppress endogenous and/or exogenous *Ctnnb1* expression (Bernkopf, Hadjihannas, & Behrens, 2015; Lustig et al., 2002). Supporting this hypothesis, we observed no difference in Ctnnb1 protein expression between control and transduced cells (Figure 1L), which suggested that endogenous Ctnnb1 was degraded or repressed while exogenous caCtnnb1 accumulated and entered the nucleus to upregualte downstream Wnt targets such as *Lef1* and *Axin2* and drive CM proliferation. Similar to caYap8SA and caCtnnb1, we also observed increased Ccnd2 gene but not protein expression in Ccnd2-transduced hiPSC-CMs (Figure 1M,N). Still, immunostaining analysis revealed that Ccnd2-transduced hiPSCMs exhibited higher nuclear abundance of the protein (Figure 1O,P), which likely contributed to the observed increase in CM cycling (Zhu et al., 2017). Overall, LV-mediated expressions of various mitogens in hiPSC-CMs appeared to induce negative feedback responses at a post-transcriptional level to limit protein overexpression, which in the case of caYap8SA prevented downstream proliferative effects.

### cahErbb2 Induces Cell Cycle Entry in NRVM Monolayers Associated with Sarcomere Disassembly

We then sought to determine whether LV-induced expression of the mitogens affected NRVMs, as these cells represent more mature CMs than the relatively immature hiPSC-CMs. NRVMs were transduced with the LVs the day of plating and analyzed by flow cytometry after two weeks of monolayer culture (Figure 2A). We found that LV transduction of cahErbb2 but not other mitogens resulted in increased EdU incorporation in NRVMs, while both Ccnd2 and cahErbb2 yielded increased H3P expression (Figure 2B,C). However, all transduced mitogens resulted in significantly decreased fraction of diploid and increased fraction of polyploid NRVMs (Figure 2D), indicating that LV-induced cycling events led to polyploidization rather than cytokinesis, in constrast to the results in hiPSC-CMs (Figure 1D). This inference was supported by the finding that compared to control LV case, the diploid fraction of EdU^+^ NRVMs was significantly decreased and polyploid fraction significantly increased in all LV-treated groups (Figure 2E). Together, these results revealed that Ccnd2 and cahErbb2 promote NRVM cell cycle entry and mitosis, while all tested mitogens induce NRVMs to become binucleated or mononuclear and polyploid, rather than completing cytokinesis. In both NRVMs and hiPSC-CMs, LV expression of cahErbb2, but not other mitogens, also induced sarcomere disassembly (Figure 2G). While, expectedely, gene expression of *Erbb2* was upregulated in cahErbb2-transduced cells, expression of sarcomeric genes was unaltered (Figure 2F), suggesting that post-transcriptional changes at a protein level were responsible for the loss of sarcomeric organization. Furthermore, increased expression of *Runx1* along with the loss of sarcomers implied that cahErbb2 induced CM dedifferentiation in addition to proliferation (D’Uva et al., 2015; Kubin et al., 2011).

**Figure 2.**
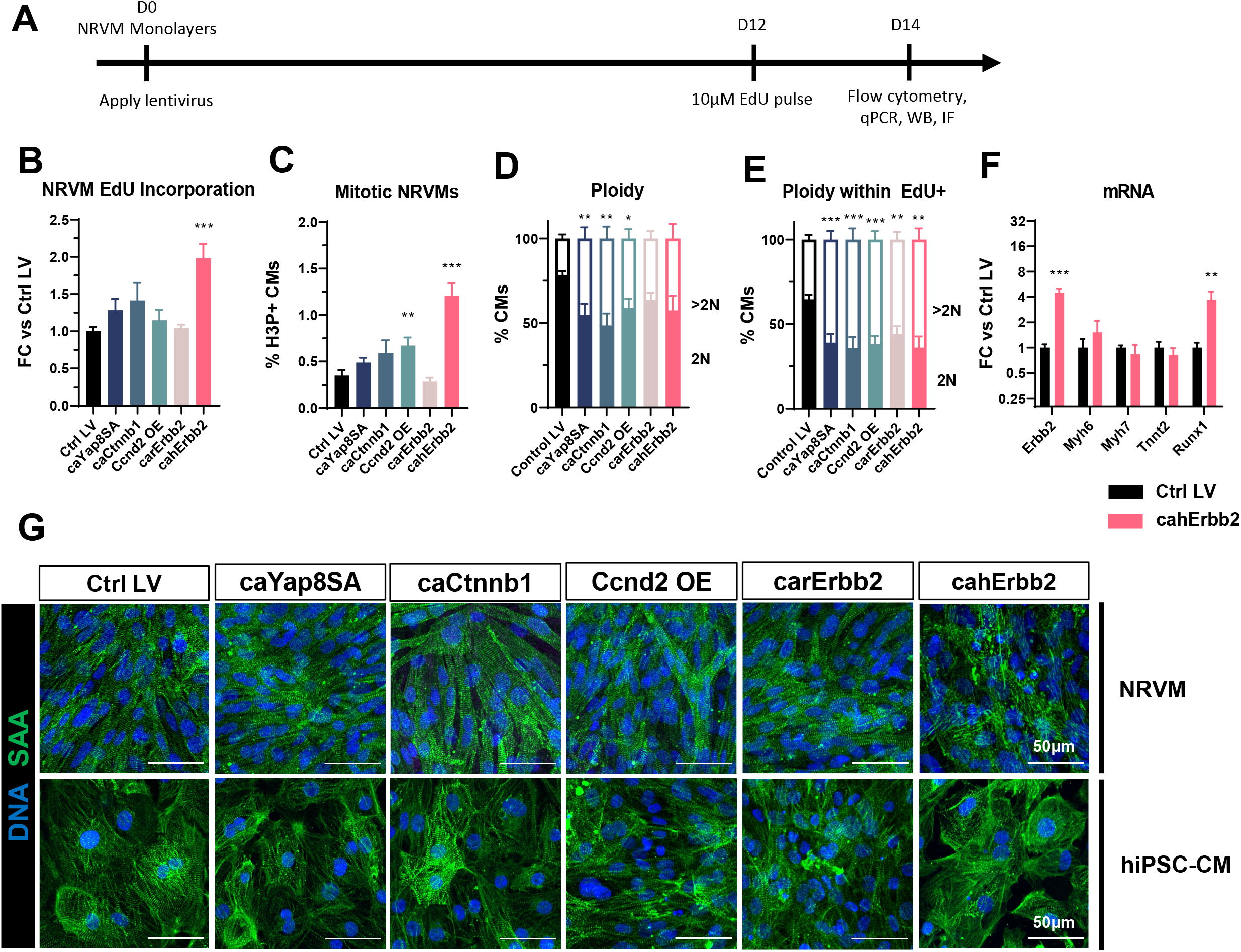
cahErbb2 Induces NRVM Cycle Entry in Monolayers and Promotes Sarcomere Disassembly in NRVMs and hiPSC-CMs. (A) Schematic of experimental design in NRVM monolayers. (B-E) Flow cytometry analysis of mCherry^+^ NRVMs showing (B) fold-change (FC) in EdU incorporation relative to control LV-treated NRVMs, (C) percentage of H3P^+^ CMs, and (D,E) percentage of 2N and >2N cells in all CMs (D) and EdU+ CMs (E). (F) Relative expression of *Erbb2*, sarcomeric genes (*Myh6*, *Myh7*, *Tnnt2*), and dedifferentiation marker *Runx1* in cahErbb2-transduced vs. control hiPSC-CMs. (G) Representative immunostaining images of sarcomeric α-actinin (SAA) showing sarcomeric structure in LV transduced NRVMs and hiPSC-CMs. Data: mean ± SEM (*p < 0.05, **p < 0.01, ***p < 0.001 vs. Ctrl LV).

### cahErbb2 Induces Cell Cycle Activation, Growth, and Contractile Deficit in NRVM Cardiobundles

We further tested whether LV delivery of mitogens affected CM proliferation and function in 3-dimensional NRVM cardiobundles, currently representing the most mature *in vitro* model of the postnatal myocardium (C. Jackman, Li, & Bursac, 2018b; C. P. Jackman et al., 2016). NRVMs were transduced at the time of cardiobundle making and assessed after two weeks of culture (Figure 3A), first ensuring that no studied structural or functional properties differed between contorl LV-transduced and untransduced tissues (Figure 3 – figure supplement 1). From cardiobundle cross-sectional stainings, EdU^+^ nuclei were observed in both vimentin^+^ cardiac fibroblasts (C. Jackman et al., 2018b; C. P. Jackman et al., 2016) predominantly residing at the tissue periphery and in F-actin^+/^ vimentin^−^ CMs (Figure 3B, top, middle). Similar to hiPSC-CM and NRVM monolayers, all studied mitogens except caYap8SA increased EdU incorporation in NRVM cardiobundles, with cahErbb2 showing the strongest effect (Figure 3C). We then assessed morphological and functional characteristics of cardiobundles and found that cahErbb2 transduction uniquely increased both the total cross-sectional area (CSA, Figure 3D) and F-actin^+^ (CM) area of cardiobundles, leading to the formation of a necrotic core (devoid of Hoechst-positive nuclei; Figure 3E), likely caused by limited diffusion of oxygen into the center of the tissue (Suvarnapathaki, Wu, Lantigua, Nguyen, & Camci-Unal, 2019). We also observed increased vimentin^+^ CSA, indicating increased fibroblast abundance (Figure 3F). By measuring the contractile force generation in cardiobundles, we found a reduced twitch amplitude in tissues transduced with cahErbb2 but not other mitogens (Figure 3G). This loss of active force generation was accompanied by significant increase in passive tension (Figure 3H), indicating increased tissue stiffness. The cahErbb2-transduced cardiobundles also showed a near-complete loss of sarcomere structure, consistent with the findings in CM monolayers (Figure 2G), while the NRVM alignment was maintained (Figure 3B, bottom). Taken together, among the LV-expressed mitogens, cahErbb2 induced the most potent proliferative effects in both rat and human CMs, which involved the sarcomere loss characteristic of CM dedifferentiation observed in carErbb2-expressing mice (D’Uva et al., 2015). Furthermore, in engineered NRVM cardiobundles, caErbb2 proliferative effects were uniquely associated with the increase in tissue size, loss of sarcomeres and contractile force, and tissue stiffening.

**Figure 3.**
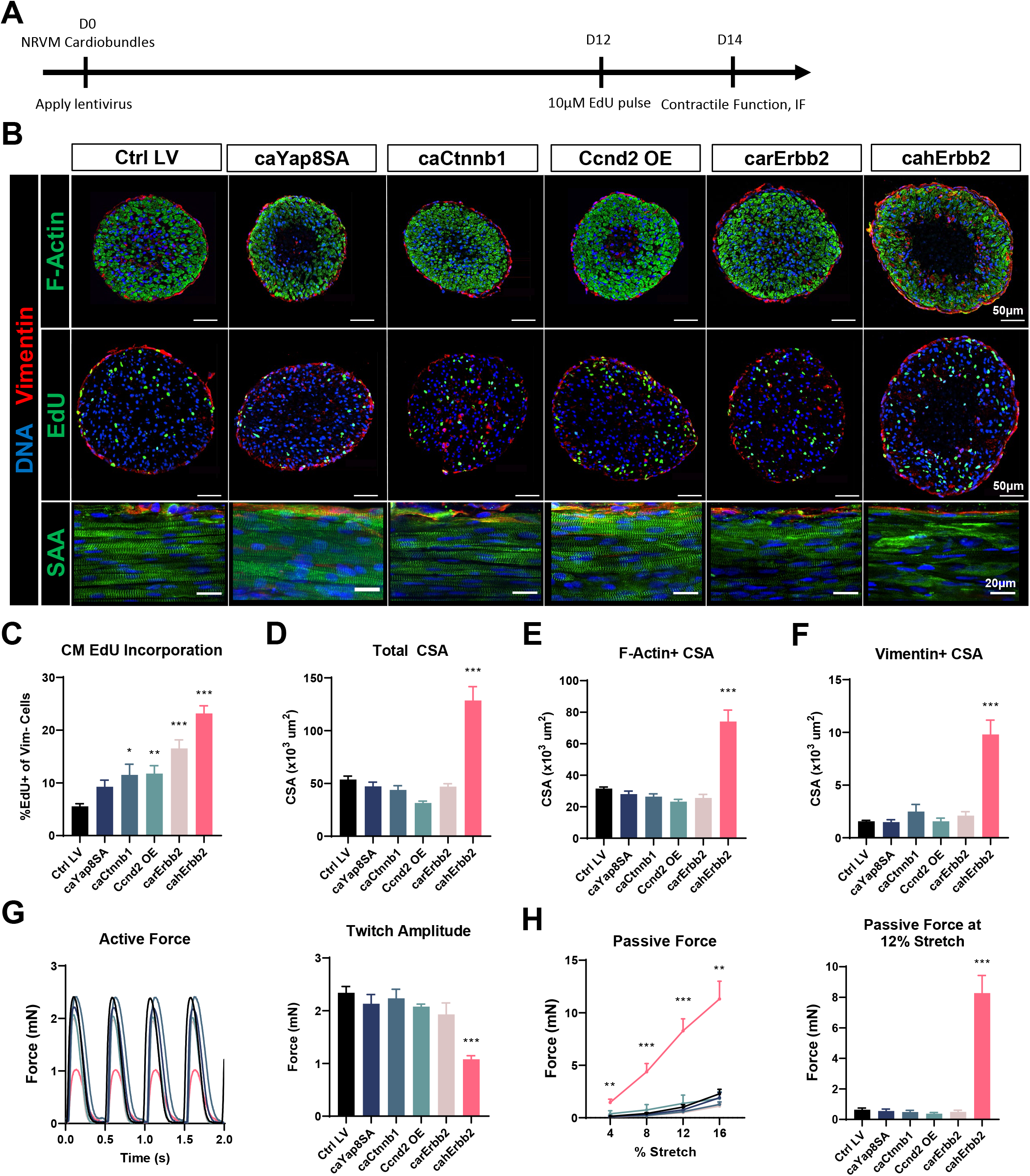
cahErbb2 Induces NRVM Cycle Entry in Cardiobundles and promotes Sarcomere Disassembly nd Contractile Dysfunction. (A) Schematic of experimental design in NRVM cardiobundles. (B) Representative immunostaining images of cardiobundle cross-sections showing morphology (top), EdU incorporation (middle), and whole-mount tissue showing sarcomere structure (bottom). (C-F) Quantification of immunostained cardiobundle cross-sections for (C) NRVM EdU incorporation and (D) total, (E) F-actin^+^, and (F) vimentin^+^ cross-sectional area (CSA). (G-H) Force analysis in LV-transduced cardiobundles showing (G) representative twitch traces and quantified maximum twitch amplitude and (H) passive force-length and force amplitude at 12% stretch. Data: mean ± SEM (*p < 0.05, **p < 0.01, ***p < 0.001 vs. Ctrl LV).

### Human but not Rat caErbb2 Activates Erk Signaling in hiPSC-CMs

We were intrigued by the finding that human but not rat caErbb2 exerted significant pro-proliferative effects in hiPSC-CMs and NRVMs, and have decided to further probe the mechanisms of cahErbb2 action. Previously, expression of rat caErbb2 in mouse CMs led to sarcomere disassembly *in vitro* and *in vivo*, while downstream Erk and Akt signaling were found to be the primary drivers of carErbb2-induced CM cell cycle activation (D’Uva et al., 2015). In hiPSC-CMs in our study, cahErbb2 but not carErbb2 expression resulted in increased pErk and Erk abundance (Figure 4A). Moreover, previously reported pAkt increase was not observed for either rat or human caErbb2 expression, with carErbb2 expression reducing total Akt and pAkt (Figure 4A). We then measured the relative gene expression of Erk downstream targets indicative of increased Erk activity (Wagle et al., 2018). Whereas cahErbb2 expression robustly increased transcription of multiple Erk targets, carErbb2 expression caused no such increase (Figure 4B). Consistent with the cahErbb2-induced Erk activation, we also observed an increase in nuclear localization of Erk protein (Figure 4C), and further confirmed Erk activation by analyzing expression of additional gene targets (Figure 4D). In contrast to Erk, Akt localization was unaffected by cahErbb2 expression (Figure 4E). Interestingly, cahErbb2 but not carErbb2 also increased the abundance of phosphorylated ribosomal protein S6 (pS6) in Western blots (Figure 4F), which was further confirmed by immunostaining (Figure 4G). pS6 is usually considered as a downstream target of mTOR, which can be activated by stimulation of Akt or Erk pathway (Roux et al., 2007; Warfel, Niederst, & Newton, 2011). Curiously, pmTOR expression was unaffected by cahErbb2 (Figure 4F) and Akt pathway was not upregulated (Figure 4A,B); thus, the human caErbb2-induced pS6 increase likely resulted from an mTOR-independent, Erk-dependent mechanism previously associated with Ser235/236 phosphorylation of pS6 (Roux et al., 2007).

**Figure 4.**
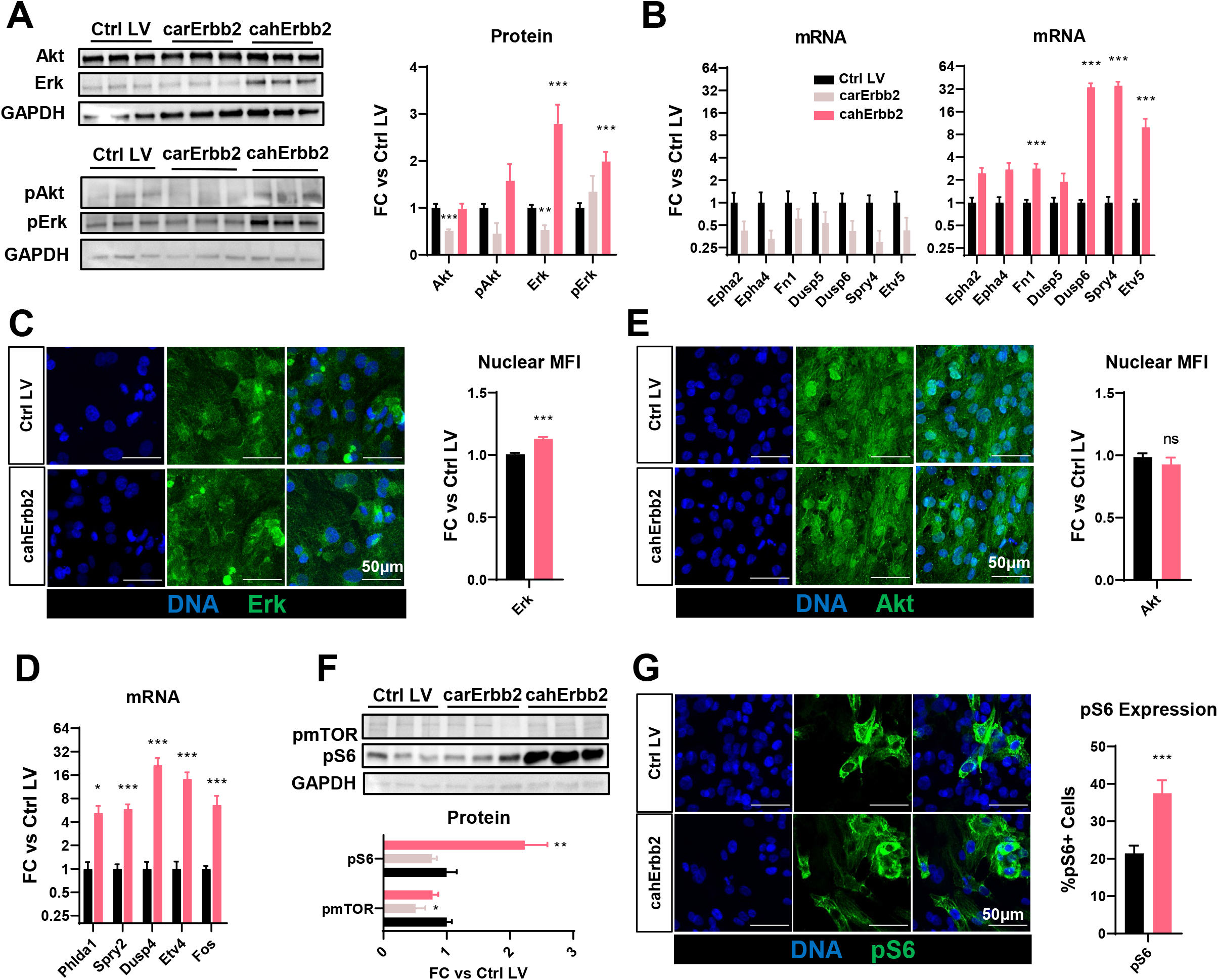
cahErbb2 but not carErbb2 Activates Erk Signaling to Drive Proliferation in CMs. (A-B) Representative Western blots and quantified relative protein (A) and Erk target gene expression (B) in carErbb2- or cahErbb2-transduced vs. control hiPSC-CM monolayers. (C) Representative immunostaining images and quantified nuclear MFI of Erk in cahErbb2-transduced vs. control hiPSC-CMs. (D) Quantified relative Erk target gene expression in cahErbb2-transduced vs. control hiPSC-CM monolayers. (E) Representative immunostaining images and quantified nuclear MFI of Akt in cahErbb2-transduced vs. control hiPSC-CMs. (F) Representative Western blots and quantified relative phosphorylated mTOR (pmTOR) and ribosomal protein S6 (pS6) expression in carErbb2- or cahErbb2-transduced vs. control hiPSC-CMs. (G) Representative immunostaining images and quantified nuclear MFI of pS6 in cahErbb2-transduced vs. control hiPSC-CMs. Data: mean ± SEM (*p < 0.05, **p < 0.01, ***p < 0.001 vs. Ctrl LV).

### Erk or Mek Inhibition Attenuates cahErbb2-induced Effects in hiPSC-CMs and NRVMs

To determine whether upregulated Erk signaling was required for the observed effects of cahErbb2 expression on hiPSC-CMs and NRVMs, we performed flow cytometry analysis in cells treated with the Mek inhibitor (Meki) PD0325901 (Mirdametinib) or the Erk inhibitor (Erki) SCH772984 applied for 48 hours before sample collection; EdU was applied during the final 24 hours to capture DNA synthesis that only occurred after adding the inhibitors (Hashimoto et al., 2014). We simultaneously measured Cc3 abundance because excessive Erk inhibition was expected to interfere with homeostatic Erk activity required to promote cell survival (Lu & Xu, 2006). As expected, in both control and cahErbb2-treated hiPSC-CMs, Meki or Erki treatment resulted in a dose-dependent decrease in EdU incorporation and a dose-dependent increase or an increasing trend in apoptotic events (Figure 5 – figure supplement 1). The increased Erk activity in cahErbb2-transduced CMs appeard to both protect against the Erki/Meki-induced apoptosis and necessicate higher inhibitor doses to maximally block EdU incorporation (Figure 5 – figure supplement 1). We then tested whether the inihibition of Erk signaling pathway can prevent cahErbb2-induced effects in NRVM cardiobundles by applying 100nM Erki or 100nM Meki between days 8 and 14 of culture (Figure 5A). We found that both inhibitors reduced CM EdU incorporation in control and cahErbb2 LV-transduced cardiobundles, but the cahErbb2-expressing tissues still showed higher rates of cycling upon inhibition compared to the control (Figure 5B, middle, 5C). Strikingly, the Erk/Mek inhibition also prevented cahErbb2-induced changes in tissue morphology, which was evident from the Erki-and Meki-reduced total, F-actin^+^, and vimentin^+^ CSA in cahErbb2-transduced but not control cardiobundles (Figure 5B,D,E,F), as well as the apperance of better-organized sarcomeric structure, especially with the Erki treatment (Figure 5B). Furthermore, the inhibitors did not affect force generation in the control cardiobundles, but partially prevented loss in active force generation (Figure 5G) and increase in passive tension (Figure 5H) in cahErbb2-transduced but not control tissues. Collectively, these results suggested that the baseline CM cycling in NRVM cardiobundles is Erk-dependent and that cahErbb2-induced morphological and functional deficits can be attenuated by Erk or Mek inhibition, further providing the evidence that Erk pathway is a major effector of cahErbb2 expression in CMs.

**Figure 5.**
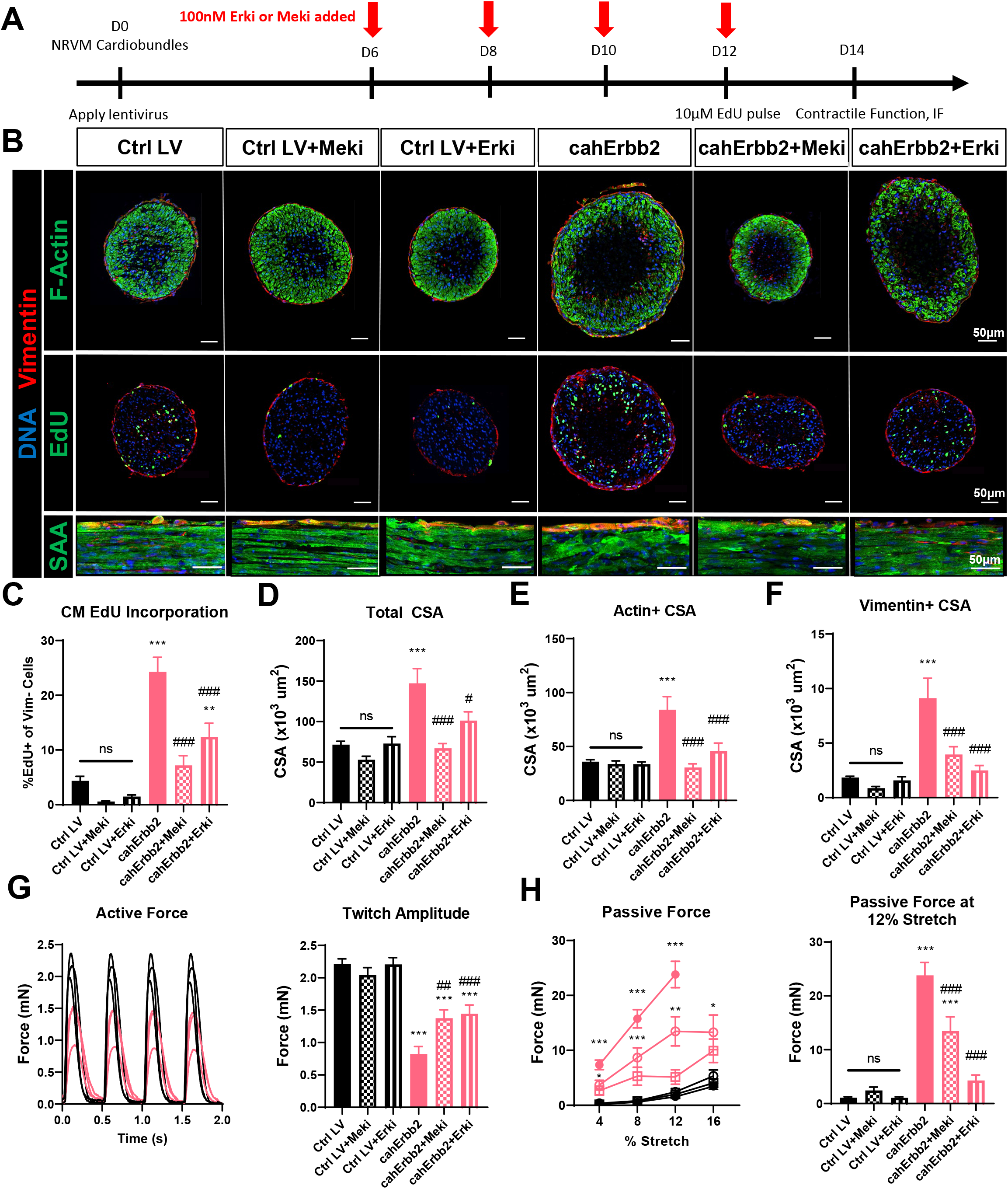
Erk or Mek Inhibition Attenuates cahErbb2-induced Effects in NRVM cardiobundles. (A) Schematic of experimental design in cahErbb2 and Ctrl LV NRVM cardiobundles. (B) Representative immunostaining images of cardiobundle cross-sections showing morphology (top), EdU incorporation (middle) and whole-mount tissue showing sarcomere structure (bottom). (C-F) Quantification of immunostained cardiobundle cross-sections for (C) NRVM EdU incorporation and (D) total, (E) F-actin^+^ (shown in Figure S5B), and (F) vimentin^+^ cross-sectional area (CSA). (G-H) Force analysis in cardiobundles showing (G) representative twitch traces and quantified maximum twitch amplitude and (H) passive force-length and force amplitude at 12% stretch. Data: mean ± SEM (*p < 0.05, **p < 0.01, ***p < 0.001 vs. Ctrl LV; ##p < 0.01, ###p < 0.001 vs. cahErbb2).

## Discussion

Development of methods to induce transient proliferation of endogenous or transplanted CMs could lead to therapeutic strategies to restore lost muscle mass and function of the injured heart (Karra & Poss, 2017). In this *in vitro* study we sought to directly compare a panel of lentivirally-delivered mitogens for the ability to induce the most potent and conserved pro-proliferative effects in CMs from different species, maturation stages, and culture environments. We found that constitutively active human Erbb2 induced a robust and more consistent proliferative response in hiPSC-CMs and NRVMs than rat caErbb2, human caCtnnb1, human Ccnd2, or human caYap8SA. Interestingly, we found apparent negative post-transcriptional feedbacks on expression of mitogen proteins following transduction with caYap8SA, caCtnnb1, or Ccnd2 LVs. This feedback appeared strongest in caYap8SA-transduced rat and human CMs, where we did not find increased EdU incorporation, while simultaneously observing a decrease in both total and active (non-phosphorylated) Yap protein. A negative feedback mechanism in the Hippo pathway mediated by Lats1/2 upregulation was previously reported when constitituvely active Yap was expressed or Sav1 was deleted in mice and cultured cells (Kim et al., 2020; Moroishi et al., 2015; Park et al., 2016). Regardless, our findings appear to contradict potent proliferative effects of constitutively active Yap mutants observed in previous studies (R. J. Mills et al., 2017; Monroe et al., 2019; von Gise et al., 2012), which may be attributable to differences in species and mechanical environment of studied CMs (Aragona et al., 2013; Benham-Pyle, Pruitt, & Nelson, 2015; Meng et al., 2018; Panciera et al., 2020; Zhao et al., 2007).

We were surprised to find that only the human but not rat ortholog of caErbb2 was able to activate Erk signaling in hCMs and NRVMs, especially considering the rat Erbb2-V663E and human Erbb2-V659E activating mutations are both located in the transmembrane domain and known to promote homo- and hetero-dimerizations of the receptor (Ou et al., 2017). Previoulsy, rat caErbb2 has been shown to drive CM dedifferentiation, proliferation, and hypertrophy when inducibly expressed in transgenic mice (Aharonov et al., 2020; D’Uva et al., 2015). In our study, the Erk activation downstream of cahErbb2 but not carErbb2 expression induced not only CM cycling, but also sarcomere disassembly acompanied by reduced contractile force generation and increased tissue stiffness and fibroblast-occupied area, which in several aspects resembled pathologic cardiac hypertrophy mediated by hyperactive Erk signaling (Gallo, Vitacolonna, Bonzano, Comoglio, & Crepaldi, 2019). While we noticed increased *Runx1* expression in cahErbb2-transduced hiPSC-CMs, we did not observe a gene expression pattern characteristic of sustained CM de-differentiation as *Myh6, Myh7,* and *Tnnt2* levels remained unchanged. Although the studied *in vitro* systems lack the complexity of an *in vivo* cardiac milieu (e.g. cellular diversity and related signaling cross-talks), it is possible that the effects of caErbb2 expression in CMs may be species or maturation-dependent. Further studies will be needed to determine whether temporally-controlled overexpression of human caErbb2 or its downstream effectors can generate comparable or more potent therapeutic effects to those of rat caErbb2 in both small and large animal studies.

## Acknowledgements

We thank A. Helfer for numerous discussions regarding the experiments performed in this manuscript, and L. Baugh for providing feedback on the initial draft of the manuscript. This work was supported by National Institutes of Health (NIH) grants U01HL134764, R01HL132389, 5T32HD040372, Foundation Leducq grant 15CVD03, and a grant from Duke Translating Duke Health Initiative. The content of the manuscript is solely the responsibility of the authors and does not necessarily represent the official views of the National Institutes of Health.

## Author Contributions

Conceptualization, N.S., S.H., R.G., and N.B.; Methodology, N.S., S.H., R.G., and N.B.; Validation, N.S. and S.D.; Investigation, N.S., S.D., and G.J.C; Data Curation, N.S. and S.D.; Writing – Original Draft, N.S., S.D., and N.B.; Writing – Review & Editing, N.S., S.D., and N.B.; Visualization, N.S. and N.B.; Supervision, N.S. and N.B.; Project Administration, N.B.; Funding Acquisition, N.B.

## Lead Contact

Further information and requests for resources and reagents should be directed to the Lead Contact, Dr. Nenad Bursac.

## Materials Availability

Plasmids generated in this study are available from the corresponding author upon reasonable request.

## Data and Code Availability

Source data for figures in the paper will be available upon request.

## Methods

### NRVM Isolation and 2D culture

All animal procedures were performed in compliance with the Institutional Animal Care and Use Committee at Duke University and the NIH Guide for the Care and Use of Laboratory Animals. Neonatal rat ventricular myocytes (NRVMs) were isolated as previously described (C. Jackman, Li, & Bursac, 2018a; C. P. Jackman et al., 2016; Li, Asfour, & Bursac, 2017). Briefly, ventricles were harvested from P2 male and female Sprague-Dawley rat pups, minced finely, and pooled before overnight trypsin incubation at 4°C. The following day, the minced ventricular tissue was subjected to several collagenase digestion and filtering steps to yield single cell suspension. Cells were pre-plated for 1 hour to remove non-myocytes and enrich the NRVM population. The non-adherent cells were resuspended in 2D cardiac medium (DMEM, 10% FBS, penicillin (5 U/ml), vitamin B12 (2 μg/ml)) and plated onto fibronectin-coated Aclar® coverslips at a density of 5×10^5^ cells per well of a 12-well plate. Twenty-four hours following plating the medium was changed to only include 5% FBS and full media changes were performed every other day.

### Cardiobundle Fabrication and 3D Culture

NRVM cardiobundles were prepared as previously described (Helfer & Bursac, 2020; C. Jackman et al., 2018a; C. P. Jackman et al., 2016). Briefly, 6.5 × 10^5^ freshly isolated NRVMs were mixed with a fibrin-based hydrogel (2.5 mg/ml fibrinogen, 1 U/ml thrombin, 10% v/v/ Matrigel) and cast in PDMS tissue molds with two 2 mm × 7 mm troughs and a porous nylon frame. The molds containing the hydrogel-cell mixture were incubated at 37°C for 45 minutes to allow the hydrogel to fully polymerize and attach to the nylon frame. Tissues were then immersed in 3D cardiac medium (Low Glucose DMEM, 10% horse serum, 1% chick embryo extract, aminocaproic acid (1 mg/ml), ascorbic acid 2-phosphate sesquimagnesium salt hydrate (50 μg/ml), penicillin (5 U/ml), vitamin B12 (2 μg/ml)). The following day, the cardiobundles on frames were carefully removed from the molds and cultured in free-floating dynamic conditions on a rocker. Full media changes of 2 mL per well were performed every other day for 14 days.

### hiPSC maintenance and CM differentiation

BJ fibroblasts from a healthy male newborn (ATCC cell line, CRL-2522) were reprogrammed episomally into hiPSCs at the Duke University iPSC Core Facility and named DU11 (Duke University clone #11) following verification of pluripotency as described previously (I. A. Shadrin, BW; Qian Y; Jackman, CP; Carlson, AL; Juhas, ME; Bursac, N, 2017). hiPSCs were maintained as feeder-free cultures on ESC-Matrigel in mTeSR Plus medium and colony-passaged as small (10–20 cells) clusters every 3 days using 0.5 mM EDTA (1:10 split ratio). All hiPSC-CM experiments were performed using DU11 hiPSCs between passages 24 - 45.

hiPSCs were differentiated into CMs via small-molecule-based modulation of Wnt signaling, as previously described (Lian et al., 2012). Briefly, DU11 hiPSCs were dissociated into single cells using Accutase and plated into ESC-Matrigel coated dishes at 4 × 10^5^/cm^2^ with 5 μM Y27632 (ROCK inhibitor). Maintenance media was changed daily prior to differentiation. To induce cardiac differentiation (day 0), cells were incubated in 6 μM CHIR99021 in RPMI-1640 with B27(−) insulin.

Exactly 48 h later, CHIR was removed and replaced with basal RPMI/B27(−) medium. On day 3, the cells were incubated in RB-containing 5 uM IWR-1, which was switched to basal RPMI/B27(−) medium on day 5. From day 7 onward, cells were cultured in RPMI/B27(+)-insulin with media changed every 2–3 days. Spontaneous contractions of hiPSC-CMs started on day 7–10 of differentiation. Differentiating CM cultures were purified via metabolic selection between days 10 and 14 (Tohyama et al., 2013), by incubation in “no glucose” medium (glucose-free RPMI supplemented with 4 mM lactate, 0.5 mg/mL recombinant human albumin, and 213 μg/mL L-ascorbic acid 2-phosphate) for 48h. At the end of the selection period, cultures were dissociated into single cells using 0.05% trypsin/EDTA followed by quenching with stop buffer (DMEM, 20% FBS, 20 μg/mL DNAse I (Millipore 260913)) and replated onto fresh Matrigel-coated dishes to remove dead cells and debris. hiPSC-CM cultures with greater than 90% cTnT positive cells as measured by flow cytometry were used for all experiments.

### Cloning of Mitogen Constructs

For generation of mitogen constructs caYap8SA, caCtnnb1, Ccnd2 OE, carErbb2, and cahErbb2, plasmid containing the gene sequence was used as PCR template for amplification of inserts prior to cloning (see Key Resources Table for plasmids). PCR primers were designed to add complementary restriction site overhangs to the gene inserts which were also present in the MHCK7-MCS-P2A-mCherry backbone used for cloning the constructs. Standard restriction cloning was used to insert the gene fragments. Sanger sequencing was performed to ensure maintenance of reading frame and correct sequence.

### Preparation of Lentivirus Vectors

Lentiviral vectors were prepared as previously described (Rao, 2018). Briefly, Hek293T cells were cultured in high glucose DMEM containing 10% FBS and 1% Penicillin/Streptomycin. Plasmids (Construct plasmid, Pax2, and VSVG) were purified using midiprep before transfection into Hek293T cells at 65-75% confluence using Jetprime transfection reagent. Medium was changed 16 hours following transfection, and medium containing virus was harvested 3-4 days following initial transfection. Virus was purified by precipitation using 3 volumes medium to 1 volume LentiX-Concentrator at 4°C overnight, then pelleted by centrifugation at 1500xg for 45 minutes at 4°C. Precipitated virus was aliquoted and stored at −80°C before use.

For experiments investigating the effects of the LVs on hiPSC-CM monolayers, cells were transduced with the LVs 17-20 days following the initiation of differentiation and were maintained for one week to allow the LVs to reach maximal expression before terminal analysis. For NRVM monolayer experiments, viral suspension was added at the time of cell plating. For NRVM cardiobundle experiments, lentiviral vectors were added to the hydrogel-cell mixture at the time of cardiobundle fabrication to yield transduction efficiency between 45-80%.

### Flow Cytometry

NRVM or hiPSC-CM monolayers were rinsed with PBS then dissociated using 0.05% Trypsin-EDTA at 37°C for 3 minutes, upon which monolayers were triturated several times to yield a single cell suspension. Trypsin was quenched with DMEM/F12 containing 20% FBS and 20μg/mL DNase I. The cell suspension was centrifuged at 300xg for 5 minutes, then resuspended in 4% PFA diluted in PBS. Cells were incubated in PFA for 10 minutes at RT, centrifuged again, then resuspended in PBS containing 5% FBS for storage at 4°C.

Cells were stained for flow cytometry after centrifugation at 300xg for 5 minutes. If EdU staining was performed, cells were incubated with the EdU flow cytometry staining cocktail as per manufacturer protocol (ThermoFisher) and incubated in the dark for 30 minutes, then washed 2X by addition of PBS followed by centrifugation. Antibody staining was performed after EdU staining. For antibody staining, cells were resuspended in FACS buffer (PBS with 0.5% BSA, 0.1% Triton-X 100, 0.02% Azide). Primary antibodies including an isotype control were diluted in FACS buffer and incubated for 1 hour on ice. Cells were washed 2X with FACS buffer before addition of secondary antibodies and Hoechst diluted in FACS buffer. Secondary antibodies were incubated for 30 minutes at RT. Samples were run on a BD Fortessa X-20.

### Immunostaining and Imaging

Cell monolayers were fixed with 4% v/v PFA at room temperature for 15 minutes, then blocked in antibody buffer (5 w/v donkey serum, 0.1% v/v Triton X-100, in PBS) for two hours at room temperature and incubated with primary antibodies for 2 hours in antibody buffer. Primary antibody dilutions are indicated in the Key Resources Table. The monolayers were washed with PBS before incubation with Alexa Fluor-conjugated secondary antibodies at 1:1000 and Hoechst at 1:200 in antibody buffer for 1 hour. Monolayer samples were mounted using Fluoromount-G™ Mounting Medium and imaged using an Andor Dragonfly spinning disk confocal microscope.

Engineered cardiobundles were fixed with 2% v/v PFA on a rocking platform at 4°C overnight. For cross-sectional analysis, the fixed tissues were suspended in OCT and flash frozen in liquid nitrogen until solidified. The frozen tissue blocks were sectioned using a cryostat (Leica) into 10μm sections. Cardiobundle cross-sections were blocked in antibody buffer for two hours at room temperature. Whole bundles for longitudinal images were blocked overnight at 4°C. All samples were incubated with primary antibodies 4°C overnight in antibody buffer. Primary antibodies were used at the indicated dilutions in the Key Resources Table. Samples were incubated with Alexa Fluor-conjugated secondary antibodies at 1:1000 and Hoechst 1:200 in antibody buffer for 2.5 hours at room temperature for cross-sections and overnight at 4°C for whole bundles. Cross-sections and un-sectioned whole bundles were mounted with hard-set mounting medium (Antifade Glass) and imaged using an Andor Dragonfly spinning disk confocal microscope.

### qPCR

RNA was extracted using RNeasy Plus Mini Kit according to the manufacturer’s instructions (Qiagen). Total RNA was converted to cDNA using iScript cDNA synthesis kit (Bio-Rad). Standard qPCR reactions with 5 ng cDNA per reaction were performed with iTaq Universal SYBR Green Supermix (Bio-Rad) in the CFX Connect Real-Time PCR Detection System. All primers used are listed in Antibody Table in Key Resources Table.

### Contractile Function

After 14 days of culture, cardiobundle force generation was measured using custom-made force measurement setup consisting of an optical force transducer and linear actuator as previously described (C. P. Jackman et al., 2016). In 37 °C Tyrode’s solution, the cardiobundle was pinned to chamber at one end and a PDMS float connected to a linear actuator controlled by Labview software at the other end. Using platinum electrodes, a 90V electrical pulse was applied for 5 ms at 2 Hz rate to induce contraction. The force measurements were performed at the ends of 4% stretch steps lasting 45 seconds, until 16% stretch was reached. Maximum twitch amplitude and passive force-length curves were calculated as previously described using custom Matlab software (Madden, Juhas, Kraus, Truskey, & Bursac, 2015).

## QUANTIFICATION AND STATISTICAL ANALYSIS

Statistical analysis was performed with GraphPad Prism software. Statistical details can be found in Supplemental Table 1. Outliers were identified and removed using GraphPad Prism 8.3.0 ROUT method, Q=1%, normality testing was done using the Shapiro-Wilk test. All experiments were carried out in multiple cell batches.

### Image Analysis

Image analysis was performed using custom CellProfiler (Carpenter et al., 2006) and FIJI (Schindelin et al., 2012) macros. Briefly, CellProfiler was used to determine nuclear number as well as vimentin^+^ and EdU^+^ cells. To quantify nuclear number and EdU^+^ nuclei, the identify primary objects function was used with global minimum cross entropy thresholding. To quantify vimentin^+^ cells, vimentin signal was smoothed using Gaussian filter, thresholded using the minimum cross entropy function, holes were removed using the remove holes function, and the watershed function was applied for segmentation. Colocalization analysis using the relate objects function between vimentin signal and nuclei as well as EdU signal and nuclei was performed to exclude proliferative fibroblasts from cardiomyocyte EdU quantification. FIJI was used to determine cardiobundle F-actin^+^ area and total cross-sectional area (CSA). For measuring F-actin+ CSA, the Huang method for auto-thresholding the F-actin signal intensity was used (Huang, 1995). For measuring total CSA, the Huang method for auto-thresholding vimentin signal intensity was used, followed by the Fill Holes function.

## Materials

| Reagent | SOURCE | IDENTIFIER |
| --- | --- | --- |
| Antibodies |  |  |
| Anti-Cardiac Troponin T antibody | Abcam | ab45932 |
| Anti-Sarcomeric Alpha Actinin antibody | Abcam | ab137346 |
| Cleaved Caspase-3 (Asp175) Antibody | Cell Signaling Technology | 9661 |
| GAPDH Antibody | Santa Cruz Biotechnology | sc-47724 |
| HER2/ErbB2 (29D8) Rabbit mAb | Cell Signaling Technology | 2165 |
| Mouse IgG1, kappa monoclonal [15-6E10A7] - Isotype Control | Abcam | ab170190 |
| Phospho-p44/42 MAPK (Erk1/2) (Thr202/Tyr204) Antibody | Cell Signaling Technology | 9101 |
| p44/42 MAPK (Erk1/2) Antibody (Total Erk) | Cell Signaling Technology | 9102 |
| Phospho-Histone H3 (Ser10) (6G3) Mouse mAb | Cell Signaling Technology | 9706 |
| Phospho-mTOR (Ser2448) Antibody | Cell Signaling Technology | 2971 |
| Phospho-p44/42 MAPK (Erk1/2) (Thr202/Tyr204) Antibody | Cell Signaling Technology | 9101 |
| Phospho-S6 Ribosomal Protein (Ser235/236) Antibody | Cell Signaling Technology | 2211 |
| Rabbit IgG, polyclonal - Isotype Control | Abcam | ab37415 |
| Recombinant Anti-active YAP1 antibody [EPR19812] | Abcam | ab205270 |
| Recombinant Anti-AKT1 (phospho S473) antibody [EP2109Y] | Abcam | ab81283 |
| Recombinant Anti-AKT1 + AKT2 + AKT3 antibody [EPR16798] | Abcam | ab179463 |
| Recombinant Anti-Cyclin D2 antibody | Abcam | ab207604 |
| Recombinant Anti-Vimentin antibody [EPR3776] - Cytoskeleton Marker | Abcam | ab92547 |
| YAP (D8H1X) XP® Rabbit mAb (Total Yap) | Cell Signaling Technology | 14074 |
| β-Catenin Antibody (Carboxy-terminal Antigen) | Cell Signaling Technology | 9587 |
| Biological Samples |  |  |
| Horse Serum (for 3D medium) |  |  |
| C3999 Chick Embryo Extract, Ultrafiltrate, Liquid (CEE) | US Biological | C3999 |
| Bovine fibrinogen | Sigma | F8630-1G |
| Thrombin from bovine plasma | Sigma | T6634-1KU |
| Chemicals, Peptides, and Recombinant Proteins |  |  |
| ICG-001 | Cayman Chemical | 16257 |
| SCH772984 | Cayman Chemical | 19166 |
| PD0325901 | Sigma | PZ0162-5MG |
| Critical Commercial Assays |  |  |
| Click-iT™ EdU Alexa Fluor™ 647 Flow Cytometry Assay Kit | Thermo Fisher | C10419 |
| Click-iT® EdU Alexa Fluor® 488 Imaging Kit | Thermo Fisher | C10337 |
| Experimental Models: Cell Lines |  |  |
| DU11 iPSC Line | Duke University Stem Cell Core |  |
| Hek293T | ATCC |  |
| Experimental Models: Organisms/Strains |  |  |
| P2 Sprague-Dawley Rat Pups | Charles River |  |
| Oligonucleotides |  |  |
| See Supplementary Table 1 for qPCR primers | This paper (or published) |  |
| Recombinant DNA |  |  |
| pLV-beta-catenin deltaN90 | pLV-beta-catenin deltaN90 was a gift from Bob Weinberg | Addgene plasmid # 36985 |
| HER2 CA (V659E) | HER2 CA (V659E) was a gift from Mien-Chie Hung | Addgene plasmid # 16259 |
| R777-E020 Hs.CCND2-nostop | R777-E020 Hs.CCND2-nostop was a gift from Dominic Esposito | Addgene plasmid # 70304 |
| pSV2 neuNT | pSV2 neuNT was a gift from Bob Weinberg | Addgene plasmid # 10918 |
| pCMV-flag YAP2 5SA | pCMV-flag YAP2 5SA was a gift from Kunliang Guan | Addgene plasmid # 27371 |
| Pax2 | psPAX2 was a gift from Didier Trono | Addgene plasmid # 12260 |
| VSVG | pMD2.G was a gift from Didier Trono | Addgene plasmid # 12259 |
| Control Lentivirus (MHCK7-mCherry) | This manuscript |  |
| Cyclin Lentivirus (MHCK7-Ccnd2-P2A-mCherry-NLS) | This manuscript |  |
| B-catenin Lentivirus (MHCK7-Ctnnb1(Δ90)-P2A-mCherry-NLS) | This manuscript |  |
| Human Erbb2 Lentivirus (MHCK7-Erbb2(V659E-P2A-mCherry)) | This manuscript |  |
| Rat Erbb2 Lentivirus (MHCK7-Erbb2(V663E-P2A-mCherry-NLS)) | This manuscript |  |
| Yap Lentivirus (MHCK7-Yap8SA-P2A-mCherry-NLS) | This manuscript |  |
| Software and Algorithms |  |  |
| CellProfiler | Mcquin et al., 2018 | www.cellprofiler.org |
| ImageJ | Schneider et al., 2012 | https://imagej.nih.gov/ij/ |
| FlowJo™ Software | Becton, Dickinson and Company; 2019 |  |
| GraphPad Prism |  | www.graphpad.com |
| Matlab |  |  |

## qPCR Primers

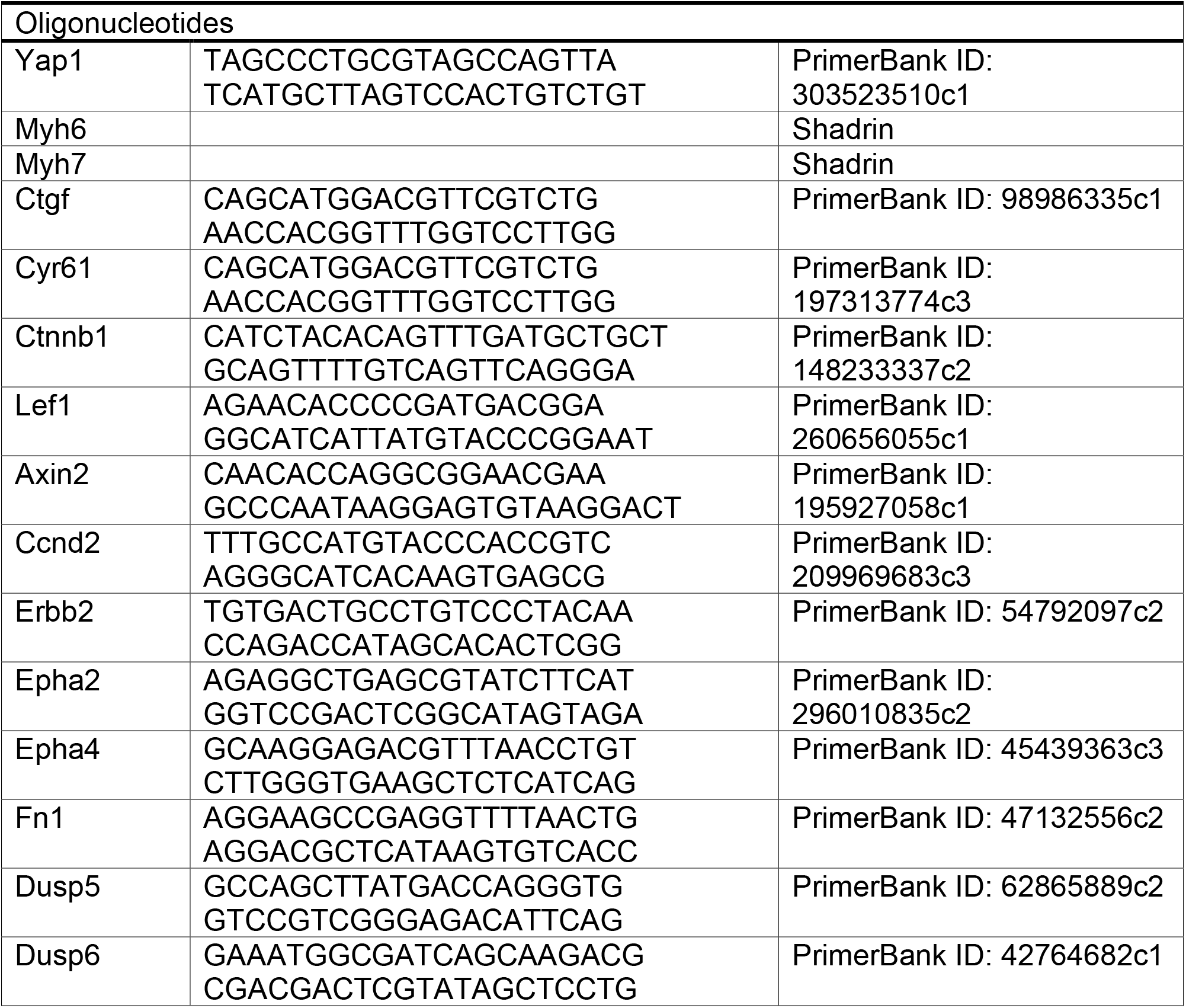
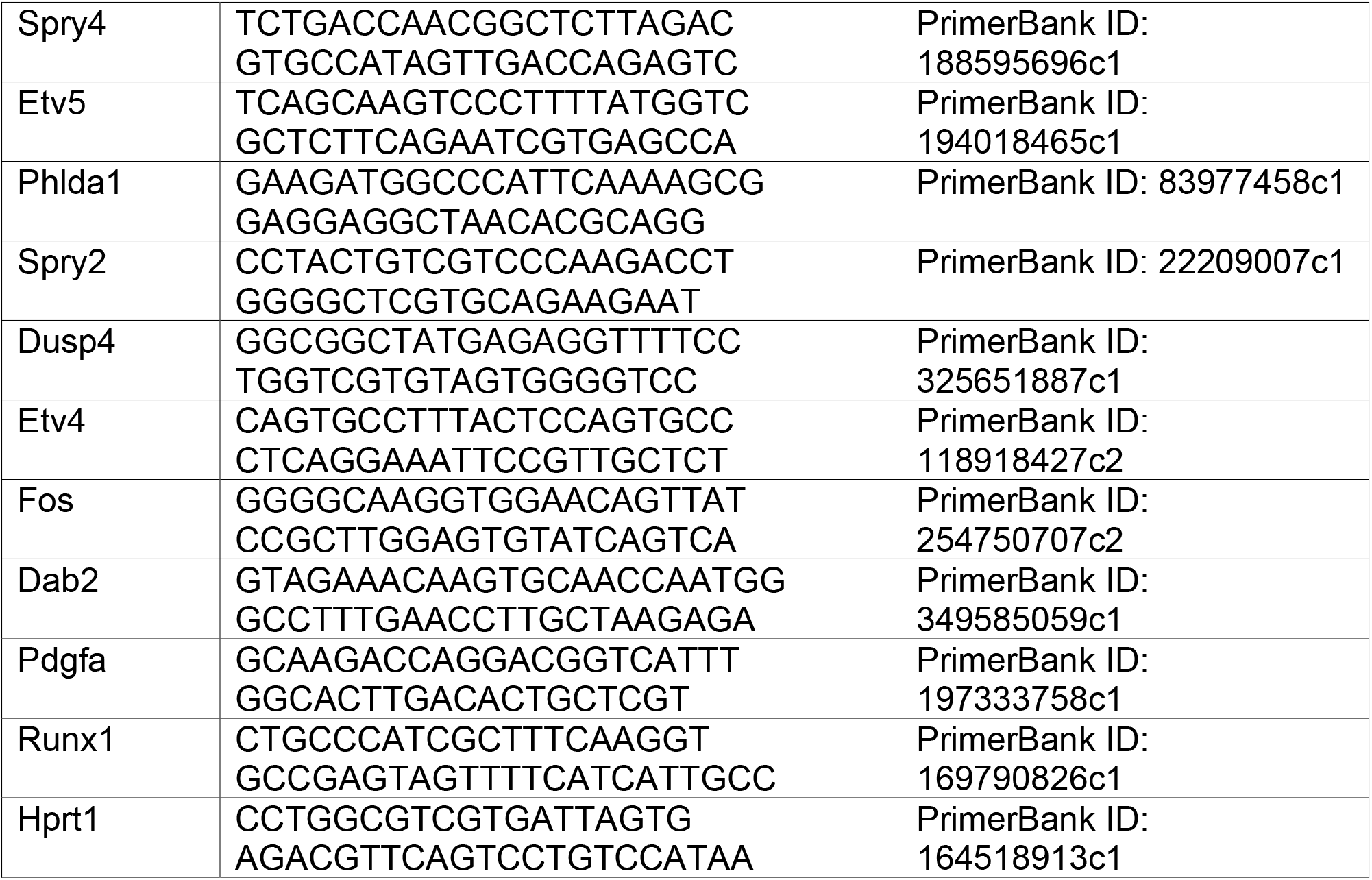

## Antibody Dilutions

### Immunofluorescence

Antibody Dilutions Immunofluorescence:
| Target | Dilution |
| --- | --- |
| SAA | 1:200 |
| Phalloidin | 1:200 |
| Ccnd2 | 1:100 |
| Yap | 1:200 |
| Active Yap | 1:500 |
| Vimentin | 1:500 |
| Akt | 1:100 |
| Erk | 1:200 |
| pS6 | 1:100 |
| Hoechst | 1:100 |

### Western Blot

Western Blot:
| Target | Dilution |
| --- | --- |
| Yap | 1:1000 |
| Active Yap | 1:1000 |
| GAPDH | 1:1000 |
| B-catenin | 1:1000 |
| Cyclin D2 | 1:1000 |
| Akt | 1:10000 |
| pAkt | 1:5000 |
| Erk | 1:1000 |
| pErk | 1:1000 |
| pmTOR | 1:1000 |
| pS6 | 1:1000 |

### Flow Cytometry

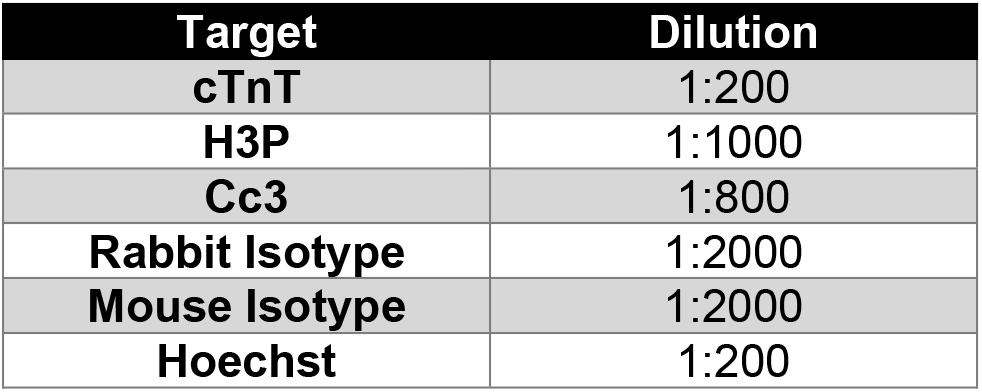

## References

Aharonov, A., Shakked, A., Umansky, K. B., Savidor, A., Genzelinakh, A., Kain, D., ... Tzahor, E. (2020). ERBB2 drives YAP activation and EMT-like processes during cardiac regeneration. Nat Cell Biol. doi:10.1038/s41556-020-00588-4

Aragona, M., Panciera, T., Manfrin, A., Giulitti, S., Michielin, F., Elvassore, N., ... Piccolo, S. (2013). A mechanical checkpoint controls multicellular growth through YAP/TAZ regulation by actin-processing factors. Cell, 154(5), 1047–1059. doi:10.1016/j.cell.2013.07.042

Benham-Pyle, B. W., Pruitt, B. L., & Nelson, W. J. (2015). Cell adhesion. Mechanical strain induces E-cadherin-dependent Yap1 and beta-catenin activation to drive cell cycle entry. Science, 348(6238), 1024–1027. doi:10.1126/science.aaa4559

Bergmann, O., Bhardwaj, R. D., Bernard, S., Zdunek, S., Barnabe-Heider, F., Walsh, S., ... Frisen, J. (2009). Evidence for cardiomyocyte renewal in humans. Science, 324(5923), 98–102. doi:10.1126/science.1164680

Bernkopf, D. B., Hadjihannas, M. V., & Behrens, J. (2015). Negative-feedback regulation of the Wnt pathway by conductin/axin2 involves insensitivity to upstream signalling. J Cell Sci, 128(1), 33–39. doi:10.1242/jcs.159145

Bersell, K., Arab, S., Haring, B., & Kuhn, B. (2009). Neuregulin1/ErbB4 signaling induces cardiomyocyte proliferation and repair of heart injury. Cell, 138(2), 257–270. doi:S0092-8674(09)00522-4 [pii] 10.1016/j.cell.2009.04.060

Buikema, J. W., Lee, S., Goodyer, W. R., Maas, R. G., Chirikian, O., Li, G., ... Wu, S. M. (2020). Wnt Activation and Reduced Cell-Cell Contact Synergistically Induce Massive Expansion of Functional Human iPSC-Derived Cardiomyocytes. Cell Stem Cell, 27(1), 50–63 e55. doi:10.1016/j.stem.2020.06.001

Byun, J., Del Re, D. P., Zhai, P., Ikeda, S., Shirakabe, A., Mizushima, W., ... Sadoshima, J. (2019). Yes-associated protein (YAP) mediates adaptive cardiac hypertrophy in response to pressure overload. J Biol Chem, 294(10), 3603–3617. doi:10.1074/jbc.RA118.006123

Carpenter, A. E., Jones, T. R., Lamprecht, M. R., Clarke, C., Kang, I. H., Friman, O., ... Sabatini, D. M. (2006). CellProfiler: image analysis software for identifying and quantifying cell phenotypes. Genome Biol, 7(10), R100. doi:10.1186/gb-2006-7-10-r100

D’Uva, G., Aharonov, A., Lauriola, M., Kain, D., Yahalom-Ronen, Y., Carvalho, S., ... Tzahor, E. (2015). ERBB2 triggers mammalian heart regeneration by promoting cardiomyocyte dedifferentiation and proliferation. Nat Cell Biol, 17(5), 627–638. doi:10.1038/ncb3149

Fan, C., Fast, V. G., Tang, Y., Zhao, M., Turner, J. F., Krishnamurthy, P., ... Zhang, J. (2019). Cardiomyocytes from CCND2-overexpressing human induced-pluripotent stem cells repopulate the myocardial scar in mice: A 6-month study. J Mol Cell Cardiol, 137, 25–33. doi:10.1016/j.yjmcc.2019.09.011

Fan, Y., Ho, B. X., Pang, J. K. S., Pek, N. M. Q., Hor, J. H., Ng, S. Y., & Soh, B. S. (2018). Wnt/beta-catenin-mediated signaling re-activates proliferation of matured cardiomyocytes. Stem Cell Res Ther, 9(1), 338. doi:10.1186/s13287-018-1086-8

Gallo, S., Vitacolonna, A., Bonzano, A., Comoglio, P., & Crepaldi, T. (2019). ERK: A Key Player in the Pathophysiology of Cardiac Hypertrophy. Int J Mol Sci, 20(9). doi:10.3390/ijms20092164

Gemberling, M., Karra, R., Dickson, A. L., & Poss, K. D. (2015). Nrg1 is an injury-induced cardiomyocyte mitogen for the endogenous heart regeneration program in zebrafish. eLife, 4. doi:10.7554/eLife.05871

Hashimoto, H., Yuasa, S., Tabata, H., Tohyama, S., Hayashiji, N., Hattori, F., ... Fukuda, K. (2014). Time-lapse imaging of cell cycle dynamics during development in living cardiomyocyte. J Mol Cell Cardiol, 72, 241–249. doi:10.1016/j.yjmcc.2014.03.020

Helfer, A., & Bursac, N. (2020). Frame-Hydrogel Methodology for Engineering Highly Functional Cardiac Tissue Constructs. Methods Mol Biol, 2158, 171–186.

Huang, L.-K. W., M-J. (1995). Image thresholding by minimizing the measure of fuzziness. Pattern Recognition, 1, 41–51.

Jackman, C., Li, H., & Bursac, N. (2018a). Long-term contractile activity and thyroid hormone supplementation produce engineered rat myocardium with adult-like structure and function. Acta Biomater, 78, 98–110. doi:10.1016/j.actbio.2018.08.003

Jackman, C., Li, H., & Bursac, N. (2018b). Long-term Contractile Activity and Thyroid Hormone Supplementation Produce Engineered Rat Myocardium with Adult-like Structure and Function. Acta Biomater. doi:10.1016/j.actbio.2018.08.003

Jackman, C. P., Carlson, A. L., & Bursac, N. (2016). Dynamic culture yields engineered myocardium with near-adult functional output. Biomaterials, 111, 66–79. doi:10.1016/j.biomaterials.2016.09.024

Karra, R., & Poss, K. D. (2017). Redirecting cardiac growth mechanisms for therapeutic regeneration. J Clin Invest, 127(2), 427–436. doi:10.1172/JCI89786

Kim, E., Kang, J. G., Kang, M. J., Park, J. H., Kim, Y. J., Kweon, T. H., ... Cho, J. W. (2020). O-GlcNAcylation on LATS2 disrupts the Hippo pathway by inhibiting its activity. Proc Natl Acad Sci U S A, 117(25), 14259–14269. doi:10.1073/pnas.1913469117

Kubin, T., Poling, J., Kostin, S., Gajawada, P., Hein, S., Rees, W., ... Braun, T. (2011). Oncostatin M is a major mediator of cardiomyocyte dedifferentiation and remodeling. Cell Stem Cell, 9(5), 420–432. doi:10.1016/j.stem.2011.08.013

Leach, J. P., & Martin, J. F. (2018). Cardiomyocyte Proliferation for Therapeutic Regeneration. Curr Cardiol Rep, 20(8), 63. doi:10.1007/s11886-018-1011-x

Li, Y., Asfour, H., & Bursac, N. (2017). Age-dependent Functional Crosstalk Between Cardiac Fibroblasts and Cardiomyocytes in a 3D Engineered Cardiac Tissue. Acta Biomater. doi:10.1016/j.actbio.2017.04.027

Lian, X., Hsiao, C., Wilson, G., Zhu, K., Hazeltine, L. B., Azarin, S. M., ... Palecek, S. P. (2012). Robust cardiomyocyte differentiation from human pluripotent stem cells via temporal modulation of canonical Wnt signaling. Proc Natl Acad Sci U S A, 109(27), E1848–1857. doi:10.1073/pnas.1200250109

Lin, Z., von Gise, A., Zhou, P., Gu, F., Ma, Q., Jiang, J., ... Pu, W. T. (2014). Cardiac-specific YAP activation improves cardiac function and survival in an experimental murine MI model. Circ Res, 115(3), 354–363. doi:10.1161/CIRCRESAHA.115.303632

Lu, Z., & Xu, S. (2006). ERK1/2 MAP kinases in cell survival and apoptosis. IUBMB Life, 58(11), 621–631. doi:10.1080/15216540600957438

Lustig, B., Jerchow, B., Sachs, M., Weiler, S., Pietsch, T., Karsten, U., ... Behrens, J. (2002). Negative feedback loop of Wnt signaling through upregulation of conductin/axin2 in colorectal and liver tumors. Mol Cell Biol, 22(4), 1184–1193. doi:10.1128/mcb.22.4.1184-1193.2002

Madden, L., Juhas, M., Kraus, W. E., Truskey, G. A., & Bursac, N. (2015). Bioengineered human myobundles mimic clinical responses of skeletal muscle to drugs. eLife, 4, e04885. doi:10.7554/eLife.04885

Meng, Z., Qiu, Y., Lin, K. C., Kumar, A., Placone, J. K., Fang, C., ... Guan, K. L. (2018). RAP2 mediates mechanoresponses of the Hippo pathway. Nature. doi:10.1038/s41586-018-0444-0

Mills, R. J., Parker, B. L., Quaife-Ryan, G. A., Voges, H. K., Needham, E. J., Bornot, A., ... Hudson, J. E. (2019). Drug Screening in Human PSC-Cardiac Organoids Identifies Pro-proliferative Compounds Acting via the Mevalonate Pathway. Cell Stem Cell. doi:10.1016/j.stem.2019.03.009

Mills, R. J., Titmarsh, D. M., Koenig, X., Parker, B. L., Ryall, J. G., Quaife-Ryan, G. A., ... Hudson, J. E. (2017). Functional screening in human cardiac organoids reveals a metabolic mechanism for cardiomyocyte cell cycle arrest. Proc Natl Acad Sci U S A, 114(40), E8372–E8381. doi:10.1073/pnas.1707316114

Mohamed, T. M. A., Ang, Y.-S., Radzinsky, E., Zhou, P., Huang, Y., Elfenbein, A., ... Srivastava, D. (2018). Regulation of Cell Cycle to Stimulate Adult Cardiomyocyte Proliferation and Cardiac Regeneration. Cell. doi:10.1016/j.cell.2018.02.014

Mollova, M., Bersell, K., Walsh, S., Savla, J., Das, L. T., Park, S. Y., ... Kuhn, B. (2013). Cardiomyocyte proliferation contributes to heart growth in young humans. Proc Natl Acad Sci U S A, 110(4), 1446–1451. doi:10.1073/pnas.1214608110

Monroe, T. O., Hill, M. C., Morikawa, Y., Leach, J. P., Heallen, T., Cao, S., ... Martin, J. F. (2019). YAP Partially Reprograms Chromatin Accessibility to Directly Induce Adult Cardiogenesis In Vivo. Dev Cell. doi:10.1016/j.devcel.2019.01.017

Moroishi, T., Park, H. W., Qin, B., Chen, Q., Meng, Z., Plouffe, S. W., ... Guan, K.-L. (2015). A YAP/TAZ-induced feedback mechanism regulates Hippo pathway homeostasis. Genes Dev, 29, 1271–1284. doi:10.1101/gad.262816

Ou, S. I., Schrock, A. B., Bocharov, E. V., Klempner, S. J., Haddad, C. K., Steinecker, G., ... Velcheti, V. (2017). HER2 Transmembrane Domain (TMD) Mutations (V659/G660) That Stabilize Homo- and Heterodimerization Are Rare Oncogenic Drivers in Lung Adenocarcinoma That Respond to Afatinib. J Thorac Oncol, 12(3), 446–457. doi:10.1016/j.jtho.2016.11.2224

Panciera, T., Citron, A., Di Biagio, D., Battilana, G., Gandin, A., Giulitti, S., ... Piccolo, S. (2020). Reprogramming normal cells into tumour precursors requires ECM stiffness and oncogene-mediated changes of cell mechanical properties. Nat Mater, 19(7), 797–806. doi:10.1038/s41563-020-0615-x

Park, G.-S., Oh, H., Kim, M., Kim, T., Johnson, R. L., Irvine, K. D., & Lim, D.-S. (2016). An evolutionarily conserved negative feedback mechanism in the Hippo pathway reflects functional difference between LATS1 and LATS2. Oncotarget, 7(17).

Rao, L. Q., Ying; Khodabukus, Alastair; Ribar, Thomas; Bursac, Nenad. (2018). Engineering human pluripotent stem cells into a functional skeletal muscle tissue. Nature Communications, volume 9,.

Roux, P. P., Shahbazian, D., Vu, H., Holz, M. K., Cohen, M. S., Taunton, J., ... Blenis, J. (2007). RAS/ERK signaling promotes site-specific ribosomal protein S6 phosphorylation via RSK and stimulates cap-dependent translation. J Biol Chem, 282(19), 14056–14064. doi:10.1074/jbc.M700906200

Salva, M. Z., Himeda, C. L., Tai, P. W., Nishiuchi, E., Gregorevic, P., Allen, J. M., ... Hauschka, S. D. (2007). Design of tissue-specific regulatory cassettes for high-level rAAV-mediated expression in skeletal and cardiac muscle. Mol Ther, 15(2), 320–329. doi:10.1038/sj.mt.6300027

Schindelin, J., Arganda-Carreras, I., Frise, E., Kaynig, V., Longair, M., Pietzsch, T., ... Cardona, A. (2012). Fiji: an open-source platform for biological-image analysis. Nat Methods, 9(7), 676–682. doi:10.1038/nmeth.2019

Shadrin, I. A., BW; Qian Y; Jackman, CP; Carlson, AL; Juhas, ME; Bursac, N. (2017). Cardiopatch platform enables maturation and scale-up of human pluripotent stem cell-derived engineered heart tissues. Nat Commun.

Shadrin, I. Y., Khodabukus, A. & Bursac, N. (2016). Striated muscle function, regeneration, and repair. Cell Mol Life Sci, 73, 4175–4202.

Suvarnapathaki, S., Wu, X., Lantigua, D., Nguyen, M. A., & Camci-Unal, G. (2019). Breathing life into engineered tissues using oxygen-releasing biomaterials. NPG Asia Materials, 11(1). doi:10.1038/s41427-019-0166-2

Tohyama, S., Hattori, F., Sano, M., Hishiki, T., Nagahata, Y., Matsuura, T., ... Fukuda, K. (2013). Distinct metabolic flow enables large-scale purification of mouse and human pluripotent stem cell-derived cardiomyocytes. Cell Stem Cell, 12(1), 127–137. doi:10.1016/j.stem.2012.09.013

Tzahor, E., & Poss, K. D. (2017). Cardiac regeneration strategies: Staying young at heart. Science, 356(6342), 1035–1039. doi:10.1126/science.aam5894

van Berlo, J. H., Kanisicak, O., Maillet, M., Vagnozzi, R. J., Karch, J., Lin, S. C., ... Molkentin, J. D. (2014). c-kit+ cells minimally contribute cardiomyocytes to the heart. Nature, 509(7500), 337–341. doi:10.1038/nature13309

von Gise, A., Lin, Z., Schlegelmilch, K., Honor, L. B., Pan, G. M., Buck, J. N., ... Pu, W. T. (2012). YAP1, the nuclear target of Hippo signaling, stimulates heart growth through cardiomyocyte proliferation but not hypertrophy. Proc Natl Acad Sci U S A, 109(7), 2394–2399. doi:10.1073/pnas.1116136109

Wagle, M. C., Kirouac, D., Klijn, C., Liu, B., Mahajan, S., Junttila, M., ... Huang, S. A. (2018). A transcriptional MAPK Pathway Activity Score (MPAS) is a clinically relevant biomarker in multiple cancer types. NPJ Precis Oncol, 2(1), 7. doi:10.1038/s41698-018-0051-4

Warfel, N. A., Niederst, M., & Newton, A. C. (2011). Disruption of the interface between the pleckstrin homology (PH) and kinase domains of Akt protein is sufficient for hydrophobic motif site phosphorylation in the absence of mTORC2. J Biol Chem, 286(45), 39122–39129. doi:10.1074/jbc.M111.278747

Zhang, D., Shadrin, I. Y., Lam, J., Xian, H. Q., Snodgrass, H. R., & Bursac, N. (2013). Tissue-engineered cardiac patch for advanced functional maturation of human ESC-derived cardiomyocytes. Biomaterials, 34(23), 5813–5820. doi:10.1016/j.biomaterials.2013.04.026

Zhao, B., Wei, X., Li, W., Udan, R. S., Yang, Q., Kim, J., ... Guan, K. L. (2007). Inactivation of YAP oncoprotein by the Hippo pathway is involved in cell contact inhibition and tissue growth control. Genes Dev, 21(21), 2747–2761. doi:10.1101/gad.1602907

Zhu, W., Zhao, M., Mattapally, S., Chen, S., & Zhang, J. (2017). CCND2 Overexpression Enhances the Regenerative Potency of Human Induced Pluripotent Stem Cell-Derived Cardiomyocytes: Remuscularization of Injured Ventricle. Circ Res. doi:10.1161/CIRCRESAHA.117.311504

